# Mapping the GALNT1 substrate landscape with versatile proteomics tools

**DOI:** 10.1101/2022.08.24.505189

**Authors:** Amir Ata Saei, Susanna L. Lundström, Hezheng Lyu, Hassan Gharibi, Weiqi Lu, Pan Fang, Xuepei Zhang, Zhaowei Meng, Jijing Wang, Massimiliano Gaetani, Ákos Végvári, Steven P. Gygi, Roman A. Zubarev

## Abstract

O-GalNAc type glycosylation is a common post-translational modification (PTM) of proteins catalyzed by polypeptide GalNAc transferases, but the substrate specificity of these transferases is poorly understood. Here we develop a strategy based on integral thermal proteome solubility profiling to identify and prioritize the protein substrates of polypeptide N-acetylgalactosaminyltransferase 1 (GALNT1). Combined with glycoprotein enrichment followed by HCD and soft EThcD gas-phase fragmentation technique, we uncover hundreds of novel GALNT1 substrates in two model human cell lines. GALNT1-mediated O-glycosylation is more common on Thr than Ser residues, with a strong preference for Pro at positions +3 and +4 in respect to O-glycosylation. These results implicate GALNT1 in potentially regulating proteins in several diverse pathways, including some unexpected processes, such as TCA cycle and DNA transcription. This study depicts a roadmap for identification of functional substrates for glycosyltransferases, facilitating fundamental insight into the role of glycosylation in homeostasis and disease.

## Introduction

O-linked glycosylation involves the attachment of saccharides to the hydroxyl group of threonine (Thr) or serine (Ser) on substrate proteins. Mucin-type O-glycosylation is initiated by a large family of 20 polypeptide GalNAc-transferases (GalNAc-Ts), which transfer activated forms of monosacharides from nucleotide sugars to protein substrates ^1^. The O-GalNAc residues are further extended by addition of different monosaccharides catalyzed by 30 or more distinct glycosyltransferases. This process is initiated in the Golgi apparatus after protein folding ^2^, giving rise to arguably the most complex and most abundant O-glycan type ^3^. GalNAc-Ts perform the first decisive step in O-glycosylation, thereby controlling the substrate pool and the sites of modification ^4^. GalNAc-type O-glycosylation has been traditionally considered to occur mostly on mucin proteins and in mucin-like domains of proteins ^5^. However, recent research has vastly increased the reported number of substrates, demonstrating that this type of modification can be found on all classes of proteins without mucin-like features being necessarily present ^6–8^.

O-Glycosylation plays an important role in homeostasis; it is present, e.g., in basement membrane and plays role in extracellular matrix organization ^4,9^ as well as cell adhesion ^10^. GalNAc-Ts have also been linked to susceptibility to disease in humans and animal models ^11^. For example, GALNT1 expression is implicated in bladder cancer ^12^, hepatocellular carcinoma ^13^, colorectal cancer ^14^, and breast cancer ^15^, while GALNT2 polymorphisms are associated with dyslipidemia ^16,17^. Subtle phenotypes are found to be associated with deficiencies of single GalNAc-T genes, and it has been suggested that there are unique functions for individual isoforms ^2^, as shown for GALNT1 and GALNT2 ^4^.

While the association of GalNAc-Ts with their substrates is important, the complexity of O-glycosylation makes it challenging to study on a proteome-wide scale. Enrichment of O-linked glycopeptides from complex biological samples is often named as a prerequisite for identification of otherwise low-abundant O-glycosylation sites ^18^. Therefore, a range of different techniques have been developed that enrich glycoproteins or glycopeptides using lectins ^19,20^, HILIC columns ^21,22^, hydrazide chemistry ^23,24^, metabolic labeling ^18,25^, extraction of O-linked glycopeptides (ExoO) using a chemoenzymatic method ^10^, and a genetically engineered cell system named “SimpleCell” ^7^. These enrichment techniques have extensively expanded the known O-glycoproteome in different biosystems and unraveled some disease mechanisms linked to O-glycosylation events.

Most of the affinity enrichment reagents are limited to sterically accessible epitopes ^26–29^. In N-glycan analysis, a universal enzyme can deglycosylate various common-core structures from proteins, but for O-linked proteins, no such enzyme is known ^30^. Mass spectrometry (MS) is a powerful technology to detect various post-translational modifications (PTMs), but MS-based approaches to study the O-glycome are challenging due to the heterogeneity of this modification, the lack of known consensus sequences and of universal or selective enrichment tools. *In cellulo* experiments are further complicated by the interplay of 20 GalNAc-Ts that control mucin-type O-glycosylation ^31,32^. The SimpleCell approach employs zinc-finger nuclease (ZFN) gene targeting to truncate the O-glycan elongation pathway in human cells, generating stable ‘SimpleCell’ lines with homogenous O-glycosylation, thus reducing the heterogeneity in O-glycan structures and aiding their identification ^7,33^. SimpleCell is a valuable technology and has delivered the first map of the human O-glycoproteome with 3000 glycosites in 12 human cell lines ^8^; however, it does not address the substrate specificity of each GalNAc-T. Later, the authors applied this technology to investigate the isoform-specific substrates of GALNT1 and GALNT2 ^4^. However, such studies are tedious and limited by the use of genetically altered cell lines with a single isoform of a given GalNAc-T. Furthermore, since isobaric/isomeric residues cannot be differentiated by MS, a small fraction of the identified sites in SimpleCell can be O-linked-N-acetylglucosamine (O-GlcNAc) rather than O-GalNAc ^34^. Due to the above challenges, limited number of substrates are known for specific glycosyltransferases and no isoform-specific substrate sequence motifs have so far been found for most enzymes in this class ^2,35,36^.

Predictions show that 83% of all proteins entering the endoplasmic reticulum (ER)–Golgi secretory pathway end up being O-glycosylated ^8^. With the 2,133 known substrate proteins for all GalNAc-Ts ^37^, we have mapped only a minor fraction of the O-glycoproteome. Therefore, development of complementary techniques avoiding the enrichment step would be highly desirable.

Recently, we devised a proteome-wide strategy called System-wide Identification and prioritization of Enzyme Substrates by Thermal Analysis (SIESTA) ^38^. Being based on thermal stability/solubility changes upon PTMs, SIESTA is orthogonal to other glycoproteomics strategies. In a regular SIESTA experiment, cell lysate aliquots are treated either with enzyme, co-substrate or their combination as well as vehicle for control. With proteomics analysis, changes in protein solubility happening due to enzymatic activity are tracked. In our proof-of-concept study, SIESTA identified several known and novel substrate candidates for selenoprotein thioredoxin reductase 1, protein kinase B (AKT1) and poly-(ADP-ribose) polymerase-10 systems ^38^. SIESTA was initially implemented using Thermal Proteome Profiling or TPP ^39^ (similar to MS-CETSA ^40^). We have recently introduced a new technique called Proteome Integral Solubility Alteration (PISA) assay with ≥10x higher throughput than that of TPP ^41^. There are several advantages in implementing SIESTA experiments with PISA readout. The instrument analysis time in a SIESTA experiment with an acceptable proteome coverage can be reduced from ~14 days for 2 replicates of TPP to ~2 days using a PISA analysis with 4 replicates. At the same time, the PISA data is less plagued by missing values. With the advent of TMTpro 16 multiplexing ^42^, a SIESTA-PISA experiment can be performed in 4 replicates within a single TMT set, with enhanced reproducibility and improved statistical robustness. Importantly, while enrichment techniques can only inform on the sites of modification, they cannot predict which O-GalNAc modification is likely to have functional significance, while the size of solubility shift in PISA can serve as an indirect indicator of the magnitude of structural change in proteins. As large structural change is more likely to affect protein function, the biggest advantage of SIESTA-PISA may be to help prioritize substrates and predict functional O-glycosylation events.

In this study, we apply SIESTA-PISA analysis to study substrates of GALNT1 (**Fig. 1**). GALNT1 uses UDP-α-D-N-acetylgalactosamine (hereafter called UDP-GalNAc) as a co-substrate to add GalNAc to the Thr and Ser residues of a target protein ^43^. Our main hypothesis is that this modification induces solubility changes in substrate proteins, rendering them amenable to identification by PISA. In parallel, we adapt an affinity purification strategy to enrich O-glycosylated proteins, combining it with proteomics analysis to identify the glycosylated sites by high resolution tandem MS (**Fig. 1**). By applying these approaches, we identify a multitude of GALNT1 substrates in two human cell lines representing colon and T lymphoblasts. We envision that our approach will facilitate discovery of the glycosyltransferase function and deciphering the role of these enzymes in various cellular processes related to health and disease.

**Fig. 1.**
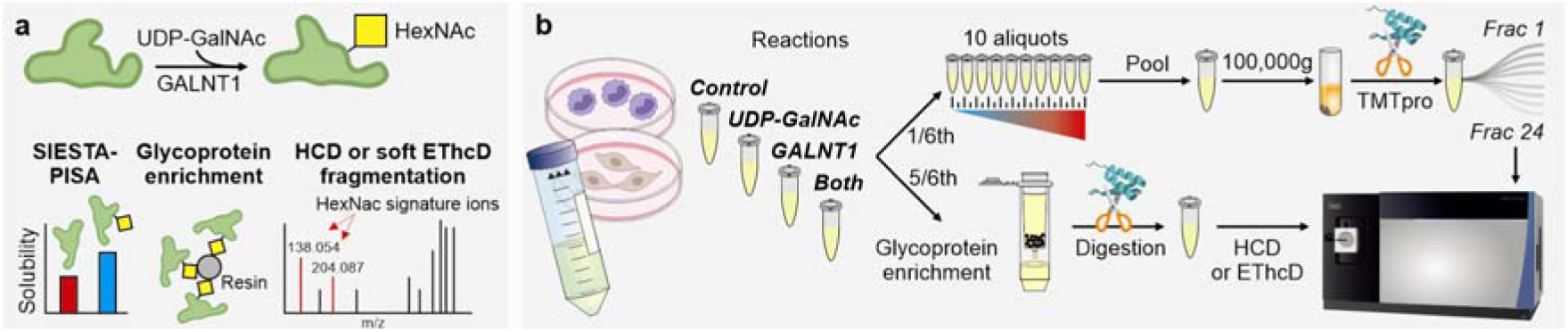
Parallel techniques for discovery and validation of GALNT1 substrates. **a,** GALNT1 employs UDP-GalNAc as a co-substrate, to add GalNAc to the Thr and Ser residues of a substrate protein, as the initial step in the O-linked oligosaccharide biosynthesis. Here we employ SIESTA with a PISA readout to discover and prioritize the GALNT1 substrates that change solubility upon O-GalNAc modification in two cell lines. In parallel, in order to verify the peptide sequence, as well as the presence and location of the glycan, glycoproteins are enriched, digested and subjected to LC-MS/MS with HCD and triggered soft EThcD fragmentation ^8^. **b,** Workflow: in SIESTA-PISA, the cell lysate is treated either with vehicle, or with UDP-GalNac, GALNT1 or both. One sixth of the total reaction volume is subjected to the PISA workflow ^41^, while the rest undergoes glycoprotein enrichment.

## Results

### Identification of GALNT1 substrates by monitoring solubility changes

Since the glycosylation machinery varies with cell and tissue type, we selected two cell lines HCT116 colorectal carcinoma cells and MOLT4 T-lymphoblasts as models. According to Human Protein Atlas, GALNT1 is expressed in both of these cell lines on an average level ^44^. Reactions took place in 400 μL of volume containing 600 μg of extracted cell protein. The cell lysate was treated with vehicle, 150 nM GALNT1, 500 μM UDP-GalNAc, or both (hereafter “Both” refers to GALNT1 + UDP-GalNAc). After 60 min of reaction time, the cell lysate was divided into 1/6^th^ and 5/6^th^ portions for the PISA and glycoprotein enrichment experiments, respectively.

In the SIESTA-PISA analysis, of the 8,643 total proteins, 7,551 proteins were quantified with 2 or more peptides (**Supplementary Data 1**). GALNT1 was detected in the two cell lines both in untreated controls and in treated samples. The endogenous concentration of GALNT1 in HEK293T cells is estimated to be around 55 nM ^45^. At 150 nM of added recombinant enzyme, the ratio of GALNT1 in treated samples vs. controls was 12.6±0.29 and 16.5±0.16 (mean±SD) for HCT116 and MOLT4 cells, respectively. Furthermore, given that the cell protein content was diluted by ~31 fold in our reactions ^38^, the recombinant GALNT1 concentration was within the physiological range.

After normalization to the total sum of intensities in each TMT channel, the normalized area under the curve (nAUC) or S_m_ for each treatment was determined by calculating the ratio of the TMT reporter ion abundances in treatment vs. control. Subsequently, the ΔS_m_ values were calculated for “Both – GALNT1” and “Both – UDP-GalNAc” (subtraction of log2 fold changes). Proteins that had a *p* value < 0.05 for “Both vs. GALNT1” and “Both vs. UDP-GalNAc” as well as a −0.05 >ΔS_m_ > +0.05 were selected as putative substrates. The scatterplots revealing known and putative substrates are shown for each cell line in **Fig. 2a-b**. The total number of unique substrates identified was 517 (90 in HCT116 and 446 in MOLT4 cells; substrates with further annotations are given in **Supplementary Data 2**). The S_m_ values for the top outliers are shown in **Fig. 2c-d**. AMER1 or APC membrane recruitment protein 1 was the top outlier. Since only 77% of the quantified proteome was shared between the two cell lines, the expected random overlap between the substrates identified in the two cell lines was 5 proteins. As we found 19 putative substrates shared between the two cell lines (**Fig. 2e**), most of these substrate candidates are true positives.

**Fig. 2.**
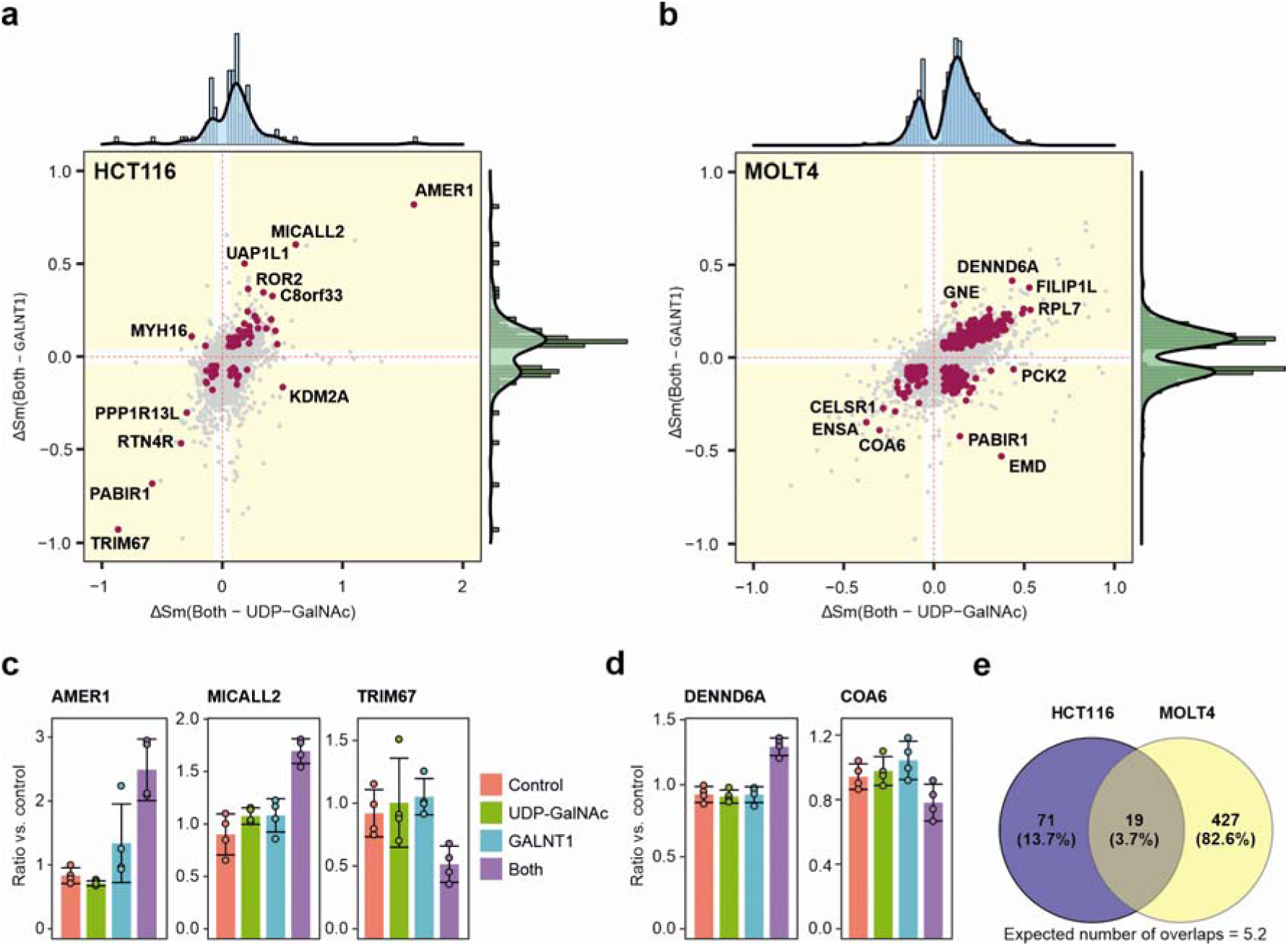
SIESTA-PISA identifies hundreds of putative GALNT1 substrates. **a-b,** Scatterplot of ΔS_m_ difference between “Both – UDP-GalNAc” and “Both – GALNT1” reveals that the solubility shifts are occurring only after simultaneous GALNT1C+CUDP-GalNAc addition compared to GALNT1 or UDP-GalNAc alone (known and putative GALNT1 substrates are shown in maroon). The plots were cropped on x and y axes to better show the distribution of the substrates. **c-d,** Representative normalized S_m_ values show the abrupt change in the solubility of putative substrates when treated with GALNT1C+CUDP-GalNAc (Both). Results are shown as mean ±SD of four independent biological replicates; two-sided Student’s t-test with unequal variance for all comparisons. **e,** The overlap between the substrates identified in the HCT116 vs. MOLT4 cells.

For better control of the false discovery rate (FDR) and higher stringency of our substrate selection, a permutation analysis of the S_m_ values in replicates was made. In total, 64,580 and 65,400 permutations (10 rounds) were performed for HCT116 and MOLT4 cells, respectively. On average less than 3 false positives were obtained in one permutation, allowing us to estimate the FDR to be ~3% for HCT116 and <0.5% for MOLT4 cells, which is significantly lower than in many glycoproteomics studies and the original SIESTA paper ^38^.

Of the identified substrates in SIESTA-PISA, 38 proteins were previously annotated in the OGP database collected from MS experiments (www.oglyp.org). These annotations were incorporated in **Supplementary Data 2**. The cellular location of 28% of the identified substrates in HCT116 cells and 34% of the substrates in MOLT4 cells was in the plasma membrane, in agreement with the well-known fact that mucin type O-glycosylation frequently occurs on membrane proteins ^46^.

### O-GalNAc tends to solubilize the substrates

While glycosylation increases the solubility of many proteins ^47^, the generality of this phenomenon has been questioned ^48^. Thus, we tested whether O-GalNAc modification mostly increases or decreases the solubility of substrate proteins. The density graphs in **Fig. 2a-b** revealed that addition of O-GalNAc tends to enhance the solubility of substrate proteins, in agreement with earlier suggestions. However, this trend is not universal, and the sign of the solubility shift is protein- and perhaps site-specific.

### PISA implicates protein-protein interactions

Furthermore, we identified proteins that interact with GALNT1 using PISA readout instead of TPP as in the original SIESTA paper ^38^. With a stringent PISA fold change cut-off of 1.5, we found 22 and 20 proteins that changed solubility significantly upon treatment with GALNT1 in HCT116 and MOLT4 cells, respectively. Four proteins were shared between the two cell lines: P4HA1, EGLN1, OGFOD2 and EIF4EBP2, while the expected number of random overlaps was around zero. With a less stringent fold change cut-off of 1.25 (**Fig. 3a-b**) we discovered 153 and 161 interacting proteins in HCT116 and MOLT4 cells (**Supplementary Data 3**). As almost all of them (98%) showed enhanced solubility upon interaction with GALNT1, true positives must compose a great majority of these proteins.

**Fig. 3.**
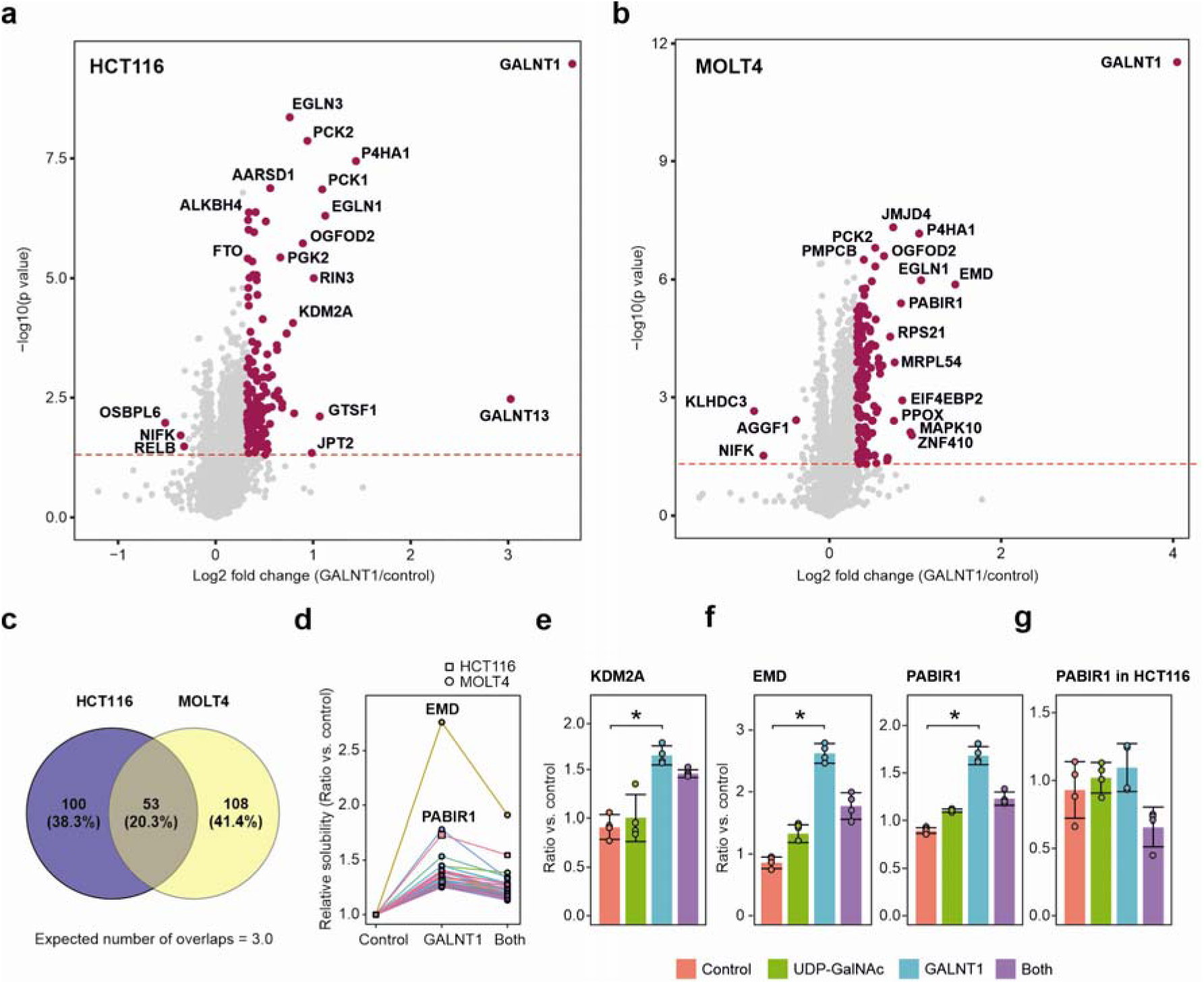
Stepwise shift in the solubility of interacting proteins which are subsequently modified. **a-b,** GALNT1 treatment alone changed the solubility of interacting proteins in different cell lines. **c,** The overlap between GALNT1 interactors in HCT116 vs. MOLT4 cells. **d,** Several interacting proteins further changed their solubility when lysate was treated with GALNT1 and UDP-GalNAc. These proteins are found in quadrant 4 in **Fig. 2a-b** and present higher solubility upon interaction with GALNT1 alone, and a loss of solubility once being modified in the presence of GALNT1 and UDP-GalNAc. Examples of putative substrates with collinear shift in solubility are given in **e-f,** KDM2A (in HCT116 cells) as well as EMD and PABIR1 (in MOLT4 cells). **g,** PABIR1 behavior is different in HCT116 cells (no enhanced solubility in the presence of GALNT1 alone, but still losing solubility when modified). Results are shown as mean ±SD of four independent biological replicates. Two-sided Student’s t-test with unequal variance (\**p*<0.05).

While there were 53 (20.3%) interacting proteins shared between the two cell lines (**Fig. 3c**) (the expected random overlap is two proteins), there was only a single protein GALNT13 listed among GALNT1 interactions in BioPlex ^49^ and BioGrid ^50^ databases. This may indicate that PISA is identifying not only strong binding events, but also transient and weak (albeit real) interactions, as we have previously postulated ^38^. Such interactions are usually not discovered in other techniques such as pulldowns where the samples are rigorously washed multiple times. These results also strengthen our previous findings on the orthogonality of thermal profiling techniques to other methods used in identification of protein interactors in lysate ^38^.

### Some substrate proteins demonstrate collinear solubility shifts

Of the 314 proteins identified by PISA as interacting with GALNT1, 45 were also found by SIESTA as GALNT1 substrates (4 for HCT116 and 41 for MOLT4 cells) (**Fig. 3d**). This is while the corresponding expected number of random overlaps is 2 and 11 proteins, respectively. Curiously, in the loading plots in **Fig. 2a-b** some proteins are situated in quadrant 4 (lower right), especially in the MOLT4 cell line. These are the proteins that increased their solubility upon interaction with GALNT1 and subsequently showed reduced solubility upon modification. Some representative examples are shown in **Fig. 3e-f**. Uniquely among all shifting proteins, PABIR1 showed a cell-dependent behavior, changing its solubility upon interaction with GALNT1 in HCT116 cells but retaining its solubility unchanged under the same conditions in MOLT4 cells. At the same time, co-treatment with GALNT1 and UDP-GalNAc decreased the solubility of PABIR1 in both cell lines, compared to lysate treated only with GALNT1 (**Fig. 2a-b** and **Fig. 3f-g**). Therefore, PABIR1 interaction with GALNT1 might be dependent on its proteoform in a given cell type.

Addition of UDP-GalNAc itself to cell lysate only shifted the solubility of few proteins, including GALE (UDP-galactose-4-epimerase) and GNE (bifunctional UDP-N-acetylglucosamine 2-epimerase/N-acetylmannosamine kinase) (**Supplementary Fig. 1**).

### SIESTA-PISA allows for monitoring modifications within the same experiment

One of the advantages of SIESTA with a PISA readout is that all the experimental conditions can be accommodated within a single TMTpro 16 set, allowing for analysis of PTMs within the same set by interrogating peptide tandem mass spectra. This is not possible in SIESTA implemented with TPP ^38^ and other TPP-dependent techniques, such as hotspot profiling ^51^. Having included O-GalNAc (HexNAc) as a variable modification in the sequence database search of SIESTA-PISA data, we detected peptides with this modification. Since the presence of multiple modifications on the same peptide is unlikely and many such identifications could be false positives, we discarded peptides containing multiple variable modifications other than O-HexNAc and N-terminal acetylation as well as the fixed Cys carbamidomethylation. Peptides carrying O-HexNAc modification were selected for further analysis. In HCT116 cells, we identified 49 peptides from 37 proteins unambiguously modified with O-HexNAc that were significantly elevated in the lysate treated with both GALNT1 and UDP-GalNAc compared to all other states (**Supplementary Fig. 2a; Supplementary Data 4**). In MOLT4 cells, we found 68 such peptides from 58 proteins (**Supplementary Fig. 2b, Supplementary Data 4**). The higher number of peptides carrying the modification in MOLT4 cells is in line with the higher number of substrates identified in this cell line compared to HCT116. Around 90% of the significantly changing modified peptides were overrepresented in the samples treated with both GALNT1 and UDP-GalNAc, validating the enrichment process. Among the identified O-glycopeptides, 26 were shared between the two cell lines, while the expected number of random overlaps was 0.4. Two and six modified peptides belonged to the substrates identified in SIESTA in HCT116 and MOLT4 cells, respectively (expected random overlap for HCT116 is 0.5 and for MOLT4 cells is 3). Also, 18 of these 69 proteins were annotated in the OGP database. We also calculated the occupancy of these sites (**Fig. 4**). Normalized modified and unmodified peptide intensities were separated, adjusted to account for missing values, and summarized to calculate HexNAc occupancy as the percentage of intensity attributed to HexNAc-modified peptides (HexNac/HexNac+Unmodified). Statistical analysis was conducted by comparing HexNAc occupancy between the “Control” and “Both” treatment groups using Welch’s t-test (Student t-test unpaired, unequal variance, two-sided), with occupancy ratios and associated *p* values calculated to assess significance (**Supplementary Data 4**).

**Fig. 4.**
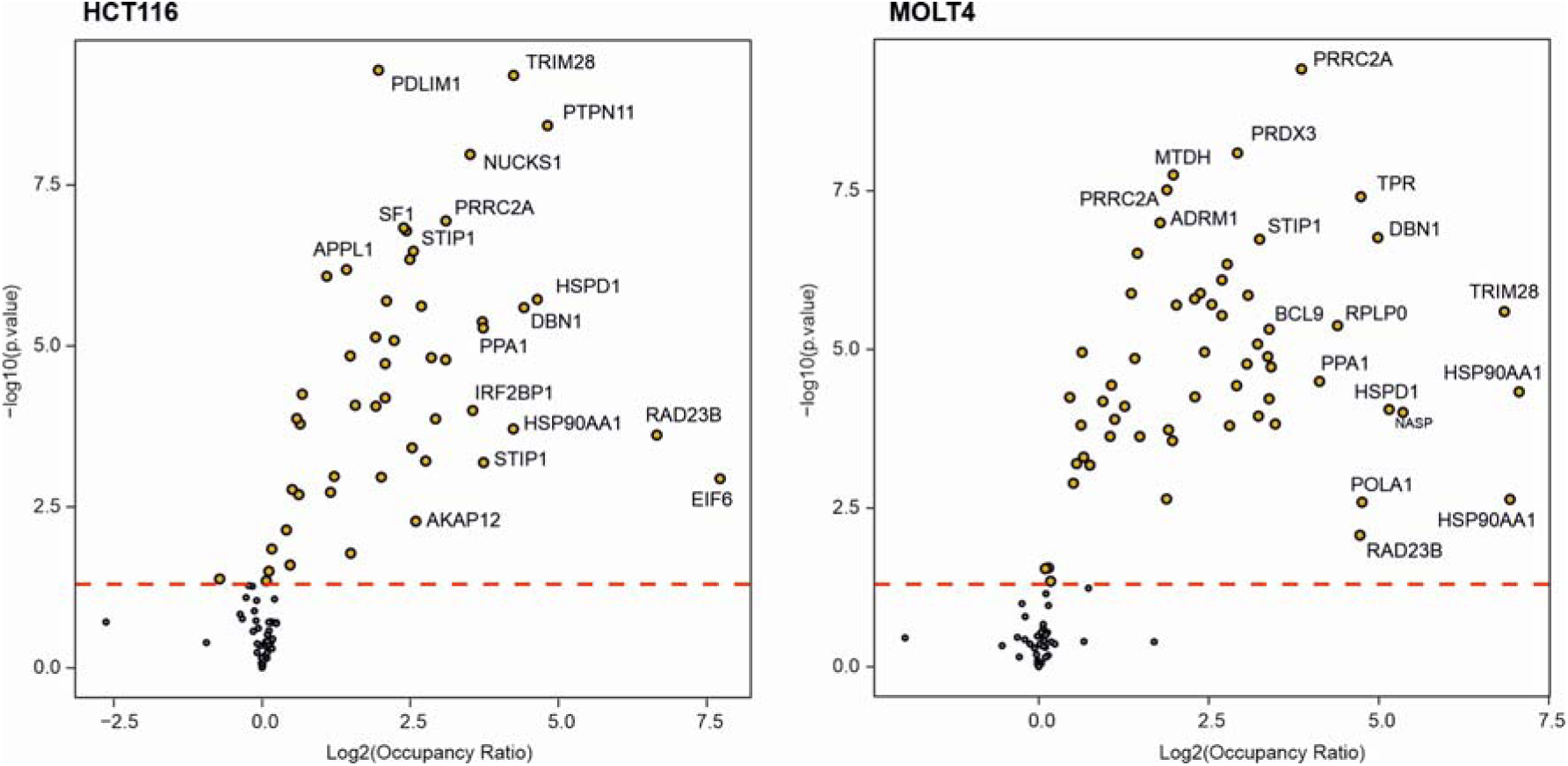
O-HexNAc modifications identified in the SIESTA-PISA experiment. Occupancy analysis showing the over-representation of O-glycopeptides a in the lysate treated with both GALNT1 and UDP-GalNAc compared to the control, validating our experiments (two-sided Welch test; four independent biological replicates).

### Glycoprotein enrichment verifies several substrates identified by SIESTA-PISA

As the samples were split prior to the thermal treatment in SIESTA-PISA analysis, 5/6^th^ of the samples were subjected to glycoprotein enrichment using GlycOCATCH kit (Genovis), as shown in **Fig. 1b**. The kit columns contain a resin that consists of agarose beads with covalently coupled inactive enzyme that only recognizes and binds to the O-glycans. Therefore, the enrichment process should be very similar to that in the EXoO approach ^10^, with the difference that here we enriched glycoproteins and not glycopeptides.

Across the two cell lines, in thus enriched samples we quantified 4,752 proteins with 2 or more peptides (**Supplementary Data 5**). In total, 86 putative substrates (44 in HCT116 and 42 in MOLT4 cells) passed a cut-off of > +0.58 for log2 fold change for Both vs. control and *p* <0.05. Of these, 6 proteins (POLR2A, B and G, as well as SNAP29, NFX1 and WEE1) were overlapping with substrates identified in SIESTA-PISA and 6 other proteins (HAGH, LMAN1 EWSR1, SNAP29, UCHL3 and SRCAP) were also annotated in the OGP database (**Supplementary Data 2**).

To identify as many O-glycan linked modification sites in peptides as possible, we combined the Proteome Discoverer ^52^ search above with a PEAKS ^53^ program database search (**Supplementary Data 6**). Applying the same exclusion criteria of peptides modified with multiple variable PTMs as above, 140 O-glycopeptides were identified in total. In **Supplementary Fig. 3**, the MS/MS spectra of representative Thr- and Ser-substituted O-glycopeptides from four proteins (RPLP0P6, CH60, RPS17 and HNRPU) are shown. Plotting the fold changes of glycopeptides in lysate treated with both GALNT1 and UDP-GalNAc vs. vehicle-treated control showed that ~94% of the significantly changing O-glycopeptides were elevated in the treated samples in both cell lines (**Fig. 5a**), hence validating the enrichment process. Furthermore, the O-glycan occupancy of the significantly changing modified peptides showed a high level of statistical significance when comparing GALNT1 + UDP-GalNAc treated lysates with control (**Fig. 5b**). Occupancy was calculated by using the log10 scaled ratio between the glycopeptide and unmodified peptide using data obtained from each corresponding peptides ion chromatogram. Each datapoint in the figure is representing the average of this log ratio in the Controls and the average in the Boths samples, respectively.

**Fig. 5.**
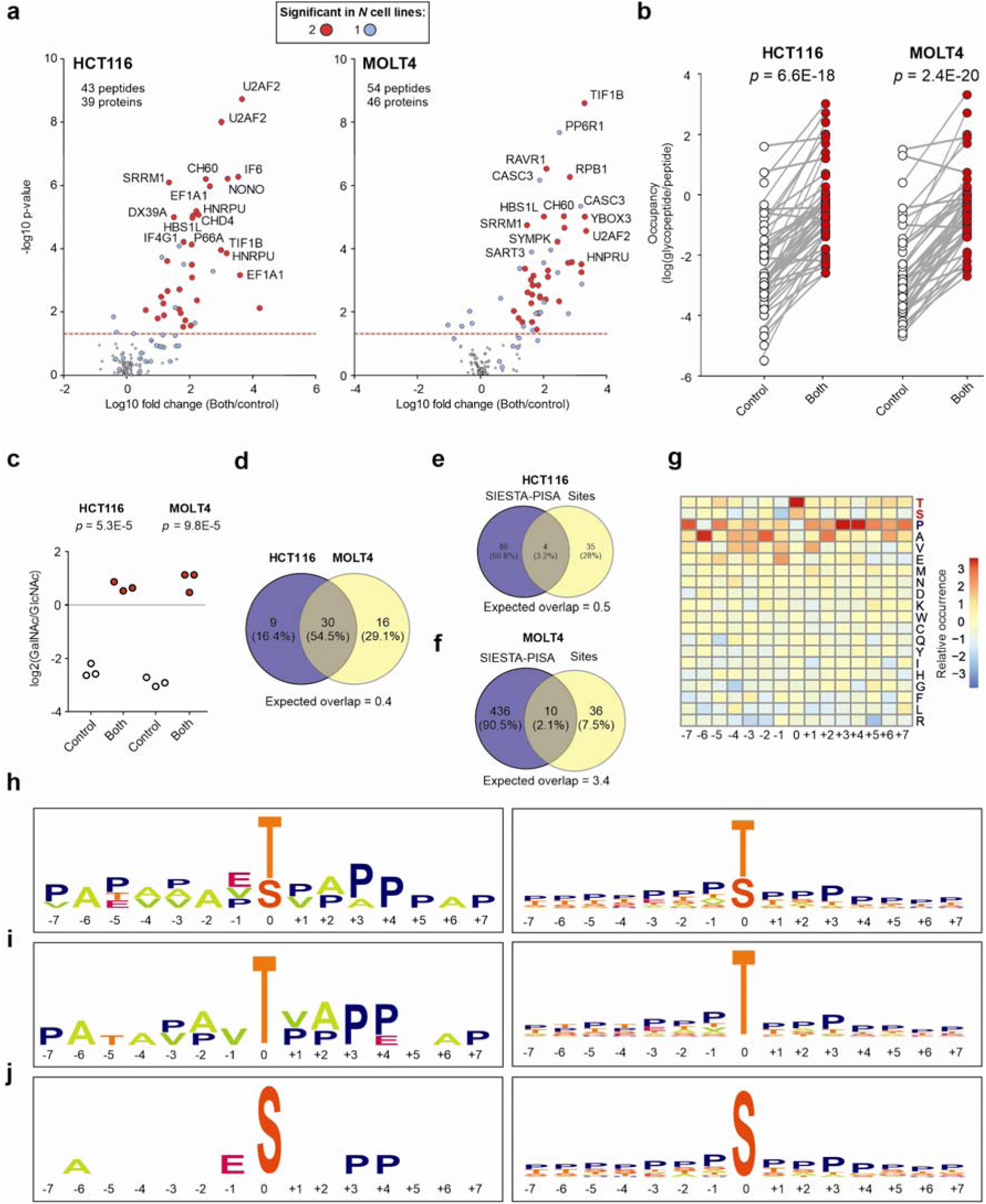
The peptides carrying O-HexNAc modification in the enrichment experiment. **a,** A majority of the glycopeptides are over-represented in the samples treated with both enzyme and UDP-GalNAc, hence validating the enrichment. Glycopeptides detected in both cell lines are shown with red circles while those identified in only one cell line are shown in blue (three independent biological replicates). **b,** The average log change between glycopeptide and corresponding unmodified peptide in the lysates treated with both GALNT1 and UDP-GalNAc vs. control that were significant in (**a**). The sample types are distinctly significant (paired two-sided Student’s t-test), thus indicating that the treated samples have a significantly higher occupancy level relatively to the controls. Note that the ratio should not be interpreted as an exact measurement of how much of the protein is modified vs unmodified. **c**, The frequency of GalNAc isomers compared to GalNAc isomers (log2[GalNAc/GlcNAc]) in the two sample types (two-sided Student’s t-test with equal variance according to F-test). **d,** The actual and expected overlap between proteins with over-represented O-HexNAc-modified peptides in the glycoprotein enrichment analysis in HCT116 vs. MOLT4 cells. **e-f,** Actual and expected overlap between substrates identified in SIESTA-PISA and proteins with elevated O-HexNAc-modified peptides in glycoprotein enrichment analysis in HCT116 and MOLT4 cells. **g,** Heatmap demonstrating the amino acid preference in the flanking regions adjacent to the modified Thr and Ser residues. **h-j,** The enriched sequence for peptides modified with O-GalNAc on Thr and Ser residues alone or together in our experiment (left panels) vs. 9,354 peptides in the OGP database (right panels).

To estimate the frequency of GalNAc isomers in the two sample types we searched the glycoprotein enriched samples using HexNAcQuest ^54^. The number of GalNAc hits in the GALNT1 + UDP-GalNAc treated lysates were 1- to 2-fold higher compared to GlcNAc and inversely the number of GlcNAc hits were 5- to 9-fold higher in the controls (**Fig. 5c** and **Supplementary Table 1)**. Note that most of the significant peptides (in **Fig. 5a**) were not characterized due to too low abundance of the diagnostic oxonium ions. However, for those that were identified, 100% were GalNAc isomers.

Of the 63 significantly enriched glycopeptides corresponding to 55 glycoproteins (**Fig. 5a**), 30 (55%) were significantly over-represented in the treated samples in both cell lines (the expected random overlap is 0.4), and 29 were specific to only one cell line (**Fig. 5d**). Only nine of these glycoproteins were annotated in the OGP database. Comparing the glycoprotein enrichment data with SIESTA-PISA, 4 (HCT116) and 10 (MOLT4) of the glycopeptides were also mapped to the putative substrate proteins identified in SIESTA-PISA (expected random overlaps are 0.5 and 3.4, respectively) (**Fig. 5e-f**). Furthermore, 10 (HCT116) and 14 (MOLT4) peptides carrying O-HexNAc in the SIESTA-PISA experiment overlapped with those identified in the glycoprotein enrichment data (the expected random overlaps are 0.3 and 0.5, respectively).

In PEAKS database search, we included the O-HexNAc-Hex modification. While 18 peptides were identified carrying O-HexNAc-Hex, only one of them (from the protein ADD1 in HCT116) was statistically significant. This indicates that in the diluted cell lysate and within the incubation period with UDP-GalNac and GALNT1, further extension of glycan units was limited.

### TxxP, TxxxP, SxxP and SxxxP are possible preferred motifs for GALNT1

While it is important to decipher and predict which Thr and Ser residues are modified with O-glycans, no clear consensus motif has been determined yet ^2,35,36^. It has been postulated that unlike N-glycosylation for which a simple consensus sequence motif exists, for O-glycosylation sequence motifs are more complicated, perhaps due to the existence of 20 GalNAc-T isoforms with different, albeit partly overlapping, substrate recognition ^2,36^. Still, some preferences for Pro, Ser, Thr, and Ala in the flanking regions around the modification site has been observed for O-glycosylation in general ^31,55^.

To identify possible motifs for GALNT1-mediated O-glycosylation, we compiled a list of significant candidate peptides carrying O-GalNAc modifications in both SIESTA and glycoprotein enrichment experiments. The list was manually curated to select the peptides with precise localization sites, resulting in 73 peptides. The curated peptides’ sequences were trimmed to equal size (15 amino acid window as done previously ^10^) by centering on modified Thr/ Ser residues and subjected to motif enrichment analysis using dagLogo ^56^. In parallel, for comparison, we performed a similar enrichment analysis for all the 9,354 sites described in the OGP database. The enrichment was performed once for peptides modified on Thr and Ser individually and once with all peptides together. The resulting heatmap for our peptides shows the preference for flanking amino acids in **Fig. 5g** and the motifs are shown in **Fig. 5h-j** for the peptides in the current study vs. peptides in the OGP database.

The sequences from the current study show an almost universal enrichment for Pro, especially in positions +3 and +4, while other positions are also frequently populated by Ala and Val. Specifically, Ala is over-represented at position −6 for peptides modified on both Thr and Ser and at position −2 only for peptides modified on Thr. Interestingly, our analysis of the OGP database (**Fig. 5h-j**) and previous research showed a generally high frequency of Pro residues around the O-glycosylation sites, with a moderate preference for +3 and −1 positions ^10,57,58^. Alanine on position −6 has also been reported to enhance the chance of glycosylation ^59^. Thus, the difference observed in flanking amino acid preferences in our data vs. OGP database can result from the fact that here we only studied GALNT1 substrates, while other analyses are made on O-glycosylation in general. Noteworthy, for GALNT2, the preferred presence of Pro at −1 and −3 has been shown ^4^. Predictions show that most O-glycosylation events are found at β turns ^60^, while proline residues are known to promote formation of β turns and β sheets ^61^.

Counting the number of modified Thr and Ser with precise localization in our curated list of modified peptides, there was a 2.2-times higher prevalence of peptides modified on Thr than Ser. Interestingly, similar analysis of the OGP database gives the same number (i.e., 2.2). Given that Thr and Ser residues comprise 5.28 and 7.26% of amino acid residues in our pulldown data from two cell lines, these results mean that Thr residues are 3.0 times more likely to be O-glycosylated. This result is in excellent agreement with a recent paper ^10^ where the authors found that O-linked glycan addition at Thr and Ser accounted for 67.6% and 22.4% of the sites, respectively.

### GALNT1 substrates map to versatile cellular compartments and pathways

We compiled a complete list of substrate candidates identified through SIESTA-PISA and glycoprotein enrichment as well as their associated MS/MS analyses in both cell lines (n = 677 proteins; **Supplementary Data 2**). We added to this table the 62 known substrates from the OGP database. In total, 71 substrate proteins identified by SIESTA-PISA were supported by at least one other technique in our experiments or literature. While larger overlap would be desirable, it should be noted that similar studies have reported ≤ 3% overlap between TPP and orthogonal techniques, such as MS-based phosphoproteomics ^38,62,63^. Among our identified substrates, several are involved in glycan machinery, such as GALNT2, RAD23B, LMAN1, ALG13 and GFPT2. Pathway analysis in Reactome found that 26 substrates are associated with asparagine N-linked glycosylation (*p*= 0.0004), interestingly indicating that perhaps both N- and O-glycosylation can occur on these substrates.

Overall, 135 proteins (20% of the total) in the compiled list were implicated by more than one analysis/database. Among these substrate candidates, there were 68% of proteins identified by pulldown MS/MS, 38% of substrate candidates identified in SIESTA MS/MS, 20% of putative substrates identified by pulldown and 13% of those identified in SIESTA-PISA.

Out of the total 677 substrate candidates in the comprehensive list, at least 156 are known to be localized to plasma membrane, in line with the known function of mucin type O-glycosylation ^46^. We further subjected the full list plus GALNT1 itself to StringDB for visualizing dense substrate hotspots and to decipher the cellular location of the substrates as well as their biological functions. The proteins mapped to several dense networks are shown in **Fig. 6**.

**Fig. 6.**
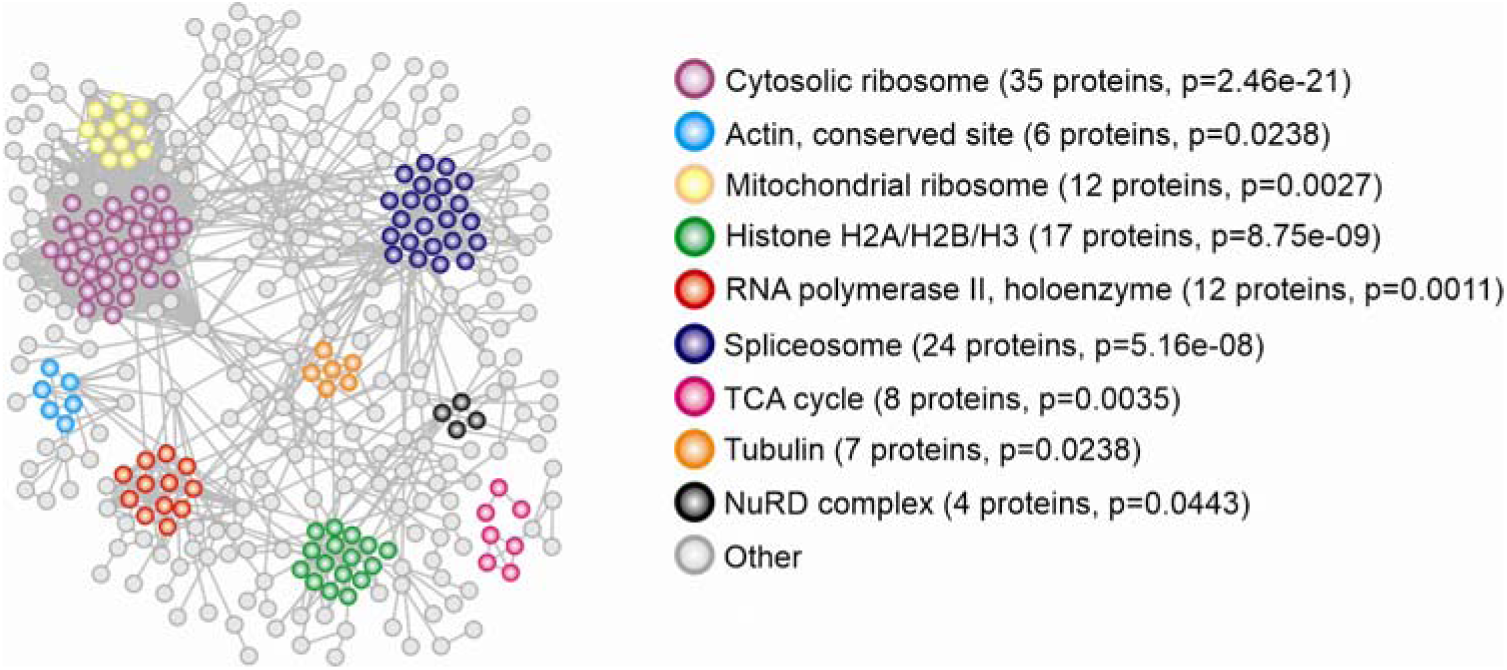
GALNT1 substrates map to different cellular compartments and complexes. The compiled list of substrates was mapped to StringDB to identify dense pathways. Only dense clusters are colored, while disconnected proteins and clusters with few members were removed for clarity.

We found that the dense enriched clusters are related to cytosolic and mitochondrial ribosome, actin, histone, RNA polymerase II, spliceosome, TCA cycle, tubulin and NuRD complex. Some other enriched pathways and processes are shown in **Table 1**.

**Table 1.** Additional processes and pathways enriched for the compiled list of GALNT1 substrates.

| Pathway | Number of proteins | P value |
| --- | --- | --- |
| SRP-dependent co-translational protein targeting to membrane | 39 | 2E-23 |
| Extracellular exosome | 146 | 1E-14 |
| Protein localization to membrane | 58 | 9E-13 |
| Cadherin binding | 36 | 1E-06 |
| Actin binding | 36 | 0.0002 |
| Intra-Golgi and retrograde Golgi-to-ER traffic | 21 | 0.0002 |
| Cell adhesion molecule binding | 40 | 0.0005 |
| Microtubule | 26 | 0.0443 |

Some of these pathways are in line with the previously known roles of O-glycosylation ^8^, such as in secretion of basement membrane components and extracellular matrix composition ^4,9^, as well as cell adhesion ^10^. The enrichment also showed various domains from InterPro ^64^, which are listed in **Supplementary Table 2**.

## Discussion

Here, parallel proteome-wide solubility- and affinity-based strategies were employed for mapping GALNT1 substrates. We report using PISA ^41^ instead of TPP as a readout for SIESTA ^38^ for the first time. This allowed us to accommodate 4 replicates of each treatment within a TMTpro 16 multiplexing set, providing robust statistics, reducing the number of missing values and allowing to investigate protein modifications within the same TMT set. Enhancing the number of replicates enabled detection of even minute changes in protein solubility. We have previously postulated that such small changes can be identified by inclusion of more replicates ^38^. With PISA, the LC-MS/MS analysis time in SIESTA is reduced from 2 weeks to 2.5 days for the same number of samples. In parallel, we adopted a modified pulldown strategy coupled to HCD or soft EThcD MS/MS fragmentation to identify substrates at the level of glycoproteins as well as O-linked glycopeptides.

Since 83% of proteins entering the ER-Golgi secretory pathway are predicted to be O-glycosylated ^8^, the current enrichment strategies, while highly informative, are not sufficient alone for mapping all substrate proteins. SIESTA-PISA provides complementary information to other available technologies, such as SimpleCell, and can be used in concert with them. Since SIESTA-PISA identifies modifications that affect protein solubility, and thus often protein conformation, these events are more likely to have functional and biological importance than the events discovered by other methods. Previous approaches compared the thermal shift of an individual modified peptide to that of the whole protein to identify “hotspot” modifications ^51,62,65^. In contrast, SIESTA-PISA compares the shift of a whole protein in the presence and absence of an enzyme and its co-substrate. Therefore, in SIESTA-PISA, the majority of protein copy numbers have to be modified for the protein to be identified as a substrate. Thus, highly ranked proteins in SIESTA-PISA are more likely to have functional relevance than in the hotspot approach that is based on peptide data.

Our results identified 677 substrate candidates for GALNT1, the majority of which are novel. The large number of putative substrates mean that GALNT1 may be involved in regulating versatile cellular processes by modifying proteins in cytosolic and mitochondrial ribosome, actin, histone, RNA polymerase II, spliceosome, TCA cycle, tubulin and NuRD complex, basement membrane, extracellular matrix and cell adhesion. Mucins and some known substrates are missing from our list. This can be ascribed to difficulties in digesting these regions because of the high density glycosylation ^8^. Also, the dominance of Ser, Thr, and Pro residues in mucin tandem repeats makes them not amenable to digestion with trypsin ^8^. Furthermore, mucins are secreted proteins and to facilitate solubility analysis we obtained the lysate without using detergents.

Interestingly, a significantly higher number of substrates were identified in MOLT4 cells compared to HCT116 cells in SIESTA-PISA. The enrichment approaches could not possibly identify such a difference without using sophisticated and costly absolute quantification techniques. Previous research has also shown substantial differences between O-glycoproteins in different cell lines ^8,66^. Future research may uncover the link between the GalNAc-T repertoire expressed in a given cell and O-glycosylation degree of its proteins.

Our results revealed that the presence of Pro at positions +3 and +4 and Ala at −6 significantly favors O-glycosylation by GALNT1 in position 0. Since comparison with general O-glycosylation events found in the OGP database showed no such trend (except for the slightly enriched +3 position), this identified sequence context may be specific to GALNT1.

The introduction of O-GalNAc on substrate proteins has a tendency toward enhancing their solubility, but this is not a general rule and a given substrate can also show a reduction in solubility upon modification. It has proven challenging to generalize the effect of PTMs on protein folding, solubility and stability ^67^. The outcome of a given modification on overall protein solubility is dictated by multiple factors, including but not limited to PTM-induced conformational distortion ^68^, conformational entropy and free energy ^69^, size and position of modification ^70^, changes in the charge state and solvent accessibility (e.g. increase in overall molecular solvent accessible surface area) ^71^. Also, given the complexity of the PTM code, the PTM cross-talk and the cumulative effect that multiple PTMs have on protein solubility ^72^, the solubility of the resulting proteoform is determined by its detailed thermodynamics ^73^. Since here we only studied the addition of O-GalNAc, a relatively small group, these results should only be cautiously extrapolated to full-size O-glycosylation events.

There are a few inherent limitations with SIESTA-PISA, some of which are also common in other techniques. Since the addition of enzyme in cell lysate can cause non-physiological modifications, orthogonal strategies must be used to validate the putative substrates as we performed in the current and previous study ^38^. While putative substrates validated that way are highly reliable, the overlap between the complementary techniques is often limited, which puts many likely substrate candidates in a gray zone. Reporting these candidates is however important, as they may be validated in a separate study by the same or another research team.

Obviously, substrates not demonstrating a shift in solubility will be missed in SIESTA-PISA ^38^. There are estimates that the solubility of only a small fraction of substrates changes significantly upon post-translational modification ^38,51,62,74^; however, the substrates undergoing larger solubility changes should be of higher biological importance. Another complication is that some acceptor Thr and Ser residues on substrates might be occupied by other common modifications, such as phosphorylation. Such modifications will prevent the modification of these sites by GALNT1, hampering their identification in cases of large occupancy. While affinity purification strategies can enrich low abundance substrates, they tend to miss weak interactions and transient binding. SIESTA can be used to fill this gap.

The SIESTA-PISA technique can provide subcellular resolution by isolating organelles under mild conditions. The use of mild detergents, such as NP40, could achieve higher representation of membrane and otherwise insoluble proteins, even though with the current approach we had an acceptable coverage of the plasma membrane proteome (1455/7011 proteins identified by Proteome Discoverer in HCT116 and 1353/7153 in MOLT4 cells mapped to plasma membrane).

We have previously shown that thermal solubility change profiling can be applied for analyzing protein-protein interactions in a cell lysate ^38^. Here, we demonstrate on the example of GALNT1 that PISA-powered discovery of protein-protein interactions provides orthogonal information to conventional techniques, such as affinity pulldowns. While some of the discovered solubility changes were shared between the two cell lines, some were cell-type specific. This can probably be attributed to the presence of diverse proteoforms in different cell lines.

Summarizing, the universality, ease, and speed of identifying enzyme-specific substrates by SIETSA-PISA can enhance our understanding of PTMs and signaling pathways in homeostasis and disease. We compiled as resource a list of GALNT1 substrate candidates and depicted a roadmap for identification of substrates for other members of the glycosyltransferase family. This high-throughput approach can be universally applied to enzymes from different classes. PISA readout allows for accommodating vehicle, co-substrate, enzyme and their combinations in four replicates within the same TMTpro 16plex set, providing robust statistics and massive reduction in the number of missing values, and yielding the desired information within a week, including sample preparation and instrumental time.

## Materials and methods

### Cell culture

Human HCT116 (ATCC, USA) cells were grown at 37°C in 5% CO_2_ in McCoy’s 5A medium supplemented with 10% FBS superior (Gibco, heat-inactivated, Fisher Scientific), 2 mM L-glutamine (Lonza BioWhittaker, Fisher Scientific) and 100 units/mL penicillin/streptomycin (Thermo). Human MOLT4 cells (ATCC, USA) were grown under the exact same conditions in RPMI 1640 Medium with GlutaMAX supplement and HEPES (Thermo). Low-number passages were used for the experiments.

### SIESTA-PISA experiments

A step-by-step protocol describing the SIESTA experiment protocol in TPP mode can be found at Protocol Exchange ^38,75^. A detailed PISA protocol can be found here ^41^. For the SIESTA-PISA experiment, cells were cultured in 175 cm^2^ flasks, and the detached by trypsin (HCT116 cells) or centrifugation (MOLT4 cells), washed twice with PBS, resuspended in PBS with complete protease inhibitor cocktail (Roche). Cells were lysed by five freeze-thaw cycles. The cell lysate was centrifuged at 10,000 *g* for 10 min and the soluble fraction was collected. The protein concentration in the lysate was measured using Pierce BCA assay (Thermo) and the lysate was equally distributed into 16 aliquots (400 μL each, 600 μg protein in each replicate) for each cell line. The aliquots were treated in 4 replicates with vehicle, 500 μM UDP-GalNAc (Sigma), 150 nM (~6 µg) active GALNT1 (ProspecBio), or with both at 37°C for 60 min.

After the reaction, for SIESTA-PISA, 100 µg of protein (in ~67 µl) from each replicate was aliquoted into 10 in 96-well plate wells and heated in an Eppendorf gradient thermocycler (Mastercycler X50s) in the temperature range of 48-59°C for 3 min. Samples were cooled for 3 min at room temperature and afterwards kept on ice. Samples from each replicate were then combined and transferred into polycarbonate thickwall tubes and centrifuged at 100,000 *g* and 4°C for 20 min.

The soluble protein fraction was transferred to new Eppendorf tubes. Protein concentration was measured in all samples using Pierce BCA Protein Assay Kit (Thermo), the volume corresponding to 25 µg of protein was transferred from each sample to new tubes and urea was added to a final concentration of 4 M. Dithiothreitol (DTT) was added to a final concentration of 10 mM and samples were incubated for 1 h at room temperature. Subsequently, iodoacetamide (IAA) was added to a final concentration of 50 mM and samples were incubated at room temperature for 1 h in the dark. Proteins were then precipitated with methanol/chloroform and then resuspended in EPPS 20 mM with 8 M urea, pH=8.5. After dilution of urea to 4 M, lysyl endopeptidase (LysC; Wako) was added at a 1:75 w/w ratio at room temperature overnight. Samples were diluted with 20 mM EPPS to the final urea concentration of 1 M, and trypsin was added at a 1:75 w/w ratio, followed by incubation for 6 h at room temperature. Acetonitrile (ACN) was added to a final concentration of 20% and TMT reagents were added 4x by weight (100 μg) to each sample, followed by incubation for 2 h at room temperature. The reaction was quenched by addition of 0.5% hydroxylamine. Samples within each replicate were combined, acidified by TFA, cleaned using Sep-Pak cartridges (Waters) and dried using DNA 120 SpeedVac Concentrator (Thermo). The pooled samples were resuspended in 20 mM ammonium hydroxide and separated into 96 fractions on an XBrigde BEH C18 2.1×150 mm column (Waters; Cat#186003023), using an Ultimate 3000 2DLC system (Thermo Scientific) over a 48 min gradient of 1-63% B (B=20 mM ammonium hydroxide in ACN) in three steps (1-23.5% B in 42 min, 23.5-54% B in 4 min and then 54-63%B in 2 min) at 200 µL min^-1^ flow. Fractions were then concatenated into 24 samples in sequential order.

### Enrichment of O-glycoproteins

500 μg in 334 μl in each replicate of the above reactions were used in parallel for enrichment of O-glycoproteins using GlycOCATCH kit (Genovis) according to the manufacturer protocols with some modifications (SialEXO and OpeRATOR enzymes were not used). The untreated and treated lysate was added to each spin column and incubated for 3 h at 25°C. After 5 washing steps, the columns were additionally washed with PBS+0.5 M urea and the bound proteins were eluted with two rounds of 8 M urea in PBS. The proteins were then digested like above and cleaned with StageTips for MS analysis in the label free mode. It is noteworthy that we also attempted to perform the enrichment in the peptide level, but the protein-level enrichment outperformed the latter analysis.

### LC-MS/MS

After drying, SIESTA-PISA samples were dissolved in buffer A (0.1% formic acid and 2% ACN in water). The samples were loaded onto a 50 cm EASY-Spray column (75 µm internal diameter, packed with PepMap C18, 2 µm beads, 100 Å pore size) connected to a nanoflow UltiMate 3000 UHPLC system (Thermo) and eluted in an organic solvent gradient increasing from 4% to 26% (B: 98% ACN, 0.1% FA, 2% H_2_O) at a flow rate of 300 nL min^-1^ over a total method of 120 min. The eluent was ionized by electrospray and mass spectra of the molecular ions were acquired with an Orbitrap Fusion Lumos tribrid mass spectrometer (Thermo Fisher Scientific) in data-dependent mode at MS1 resolution of 120,000 and MS2 resolution of 60,000, in the m/z range from 375 to 1500. Peptide fragmentation was performed via higher-energy collision dissociation (HCD) with energy set at 35% NCE and MS2 isolation width at 1.6 Th.

The glycoprotein enrichment samples were loaded like above and eluted in an organic solvent gradient increasing from 4% to 26% (B: 98% ACN, 0.1% FA, 2% H_2_O) at a flow rate of 300 nL min^-1^ over a total method of 140 min. The eluted peptides were subjected to electrospray ionization and analyzed on an Orbitrap Fusion Lumos tribrid mass spectrometer. The survey mass spectrum was acquired at the resolution of 120,000 in the m/z range of 350-1800. The first MS/MS event data were obtained with a HCD at 28% excitation for ions isolated in the quadrupole with a *m/z* width of 2 at a resolution of 30,000. Mass trigger filters targeting O-GalNAc (*m/z* 204.0867), O-GalNAc fragment (*m/z* 138.0545) and O-GalNAc Hex (*m/z* 366.1396) ions were used to initiate a second electron-transfer dissociation (ETD) with a supplementary HCD activation (EThcD) MS/MS event using ETD MS/MS with HCD supplementary activation at 15% collision energy and with a 30,000 resolution.

### Data processing

Thermo Xcalibur 4.0 was used to control and process the LC-MS data. The raw LC-MS data for SIESTA-PISA were analyzed by Proteome Discoverer version 2.5.0.400 with TMTpro 16plex as an isobaric labeling mass tag. The Sequest search engine matched MS/MS data against the UniProt complete proteome database. Cysteine carbamidomethylation was set as a fixed modification, while oxidation (M) and O-GalNAc (S and T) were selected as variable modifications. Trypsin/P was selected as enzyme specificity. No more than two missed cleavages were allowed. A 5% FDR was used as a filter at both protein and peptide levels. For all other parameters, the default settings were used.

Glycopeptide search analysis on O-linked glycan enriched samples were performed using both Proteome Discoverer and PEAKS. The Proteome Discoverer was similar to the above except that label-free quantification was used. For PEAKS, MS/MS spectra were searched against a human reference proteome with two missed cleavages as well as 10 ppm and 50 ppm mass tolerances for precursor and fragment peaks, respectively. Carbamidomethylation of cysteine was set as a fixed modification. N-Acetylhexosamine (O-GalNAc) on S and T as well as the same modification but with an additional hexose linked to the O-GalNAc moiety (Hex1O-GalNAc 1) was set as variable modifications. Search results were further validated by only including peptides with an FDR below 1% and an A score above 20, respectively.

Glycopeptide ion abundances (*m/z*, *z* and retention times) were merged, and glycopeptides were quantified in a label-free manner via in-house Quanti software ^76^, similar to what has previously been described ^77^. The program was set to detect and quantify the precursor ions of glycopeptides identified by PEAKS and Proteome Discoverer using their accurate monoisotopic masses (within <10 ppm from the theoretical values) and within ±1 min range from the expected retention times.

O-GalNAc vs O-GlcNAc frequency analysis was performed on the glycoprotein enriched samples using HexNAcQuest, https://oglcnac.shinyapps.io/o-glcnac-quest/ ^54^.

### Network mapping

For pathway analyses, STRING-DB version 11.5 protein network analysis tool ^78^ was used with default parameters.

### Statistical Analysis

Most of the data analysis was performed using R project versions 4.0 and above. Before data analysis, contaminant proteins and those identified with less than two peptides were removed from the dataset. In SIESTA-PISA, proteins with missing values were removed, and protein intensities were normalized to the sum of each channel. Then, log2 fold changes (AUC) were calculated for “UDP-GalNAc”, “GALNT1” and “Both” conditions vs. control. ΔAUCs were calculated by subtracting AUC of “UDP-GalNAc” from “Both” and “GALNT1” from Both. *P* values were calculated using two-sided Student’s t-test between normalized intensities of the 4 replicates. Proteins that had a *p* value < 0.05 for “Both vs. GALNT1” and “Both vs. UDP-GalNAc” as well as −0.05 >ΔS_m_ > +0.05 were selected as putative substrates. To assess the reproducibility of the significance cutoff used in SIESTA-PISA, a permutation analysis of the replicates was performed as described above and previously ^38,74^, validating the robustness of our discoveries.

For analysis of the pulldown results, missing values were replaced by 1/10^th^ of the minimum intensity for any given protein in each cell line and proteins LFQ values were normalized by the total abundance in each sample. Enriched proteins with a *p* value <0.05 in two-sided Student’s t-test and a log2 fold change ratio of > +0.585 were considered as substrates.

For analysis of O-GalNAc modifications in SIESTA-PISA and glycoprotein enrichment, peptide intensities were normalized to the total intensity in each channel/sample. The missing values in the whole dataset were assigned to 1/10^th^ of the minimum intensity in a given sample in a given cell line, to ensure compatibility with downstream analysis. Only peptides with 1-2 O-GalNAc modifications, those modified with carbamidomethyl on Cys and acetylation on N-term were analyzed, and peptides with multiple modifications were discarded.

For calculating the expected number of overlaps between two techniques/approaches, the size of the common dataset (number of shared proteins obtained using the two techniques) was first calculated. Then the product of the number of hits in each technique found in this common dataset was divided by the total number of proteins in the common dataset.

## Data availability

The authors declare that all data supporting the findings of this study are available within the paper and its supplementary information files. All relevant data are available from the corresponding authors (A.A.S. and R.A.Z.). The mass spectrometry data that support the findings of this study have been deposited in ProteomeXchange Consortium (https://www.ebi.ac.uk/pride/) via the PRIDE partner repository ^79^ with the dataset identifiers PXD035972 for SIESTA-PISA, PXD035991 for glycoprotein enrichment experiments.

## Acknowledgements

The research is funded by grants from the Cancerfonden (19 0558 Pj) and KAW (2019.0059) awarded to R.Z.; A.A.S. was supported by the Swedish Research Council (grant 2020-00687) and the Swedish Society of Medicine (grant SLS-961262, 1086 Stiftelsen Albert Nilssons forskningsfond) and Karolinska Institute funds (grant FS-2020:0007).

## Author contributions

Conceptualization, AAS, RAZ and SPG; project organization, resources, and funding acquisition, AAS and RAZ; methodology and experiment design, AAS, SLL, SPG and RAZ; MS experiments, AAS, HL, XZ, ZM, and MG; data analysis and visualization, AAS, SLL, HG, PF, JW, WL and AV; writing—original draft, AAS, SLL, SPG and RAZ; writing, review & editing, all co-authors.

## Competing interests

The authors declare no competing interests.

## Notes

### Competing Interest Statement

The authors have declared no competing interest.

### Summary of Updates

Figures 4 and 5 (panel c) revised. New supplementary figures 2 and 3 added. New supplementary data 7 added, and supplemental data updated.

## References

1. Varki, A., et al. Essentials of glycobiology, third edition. Cold Spring Harbor Laboratory Press (2017).

2. Bennett, E. P. et al. Control of mucin-type O-glycosylation: A classification of the polypeptide GalNAc-transferase gene family. Glycobiology at 10.1093/glycob/cwr182 (2012).

3. Joshi, H. J., et al. Protein O-GalNAc Glycosylation: The Most Complex and Differentially Regulated PTM. in Glycoscience: Biology and Medicine (2014). doi:10.1007/978-4-431-54836-2_63-1.

4. Schjoldager, K. T. et al. Deconstruction of OCglycosylation—Gal NA cCT isoforms direct distinct subsets of the OCglycoproteome. EMBO Rep. (2015) doi:10.15252/embr.201540796.

5. Jentoft, N. Why are proteins O-glycosylated? Trends Biochem. Sci. (1990) doi:10.1016/0968-0004(90)90014-3.

6. Schjoldager, K. T. B. G. et al. O-glycosylation modulates proprotein convertase activation of angiopoietin-like protein 3: Possible role of polypeptide GalNAc-transferase-2 in regulation of concentrations of plasma lipids. J. Biol. Chem. (2010) doi:10.1074/jbc.M110.156950.

7. Steentoft, C. et al. Mining the O-glycoproteome using zinc-finger nuclease-glycoengineered SimpleCell lines. Nat. Methods (2011) doi:10.1038/nmeth.1731.

8. Steentoft, C. et al. Precision mapping of the human O-GalNAc glycoproteome through SimpleCell technology. EMBO J. (2013) doi:10.1038/emboj.2013.79.

9. Tian, E., Hoffman, M. P. & Ten Hagen, K. G. O-glycosylation modulates integrin and FGF signalling by influencing the secretion of basement membrane components. Nat. Commun. (2012) doi:10.1038/ncomms1874.

10. Yang, W., Ao, M., Hu, Y., Li, Q. K. & Zhang, H. Mapping the OCglycoproteome using siteCspecific extraction of OClinked glycopeptides (EXoO). Mol. Syst. Biol. (2018) doi:10.15252/msb.20188486.

11. Tabak, L. A. The role of mucin-type O-glycans in eukaryotic development. Seminars in Cell and Developmental Biology at 10.1016/j.semcdb.2010.02.001 (2010).

12. Li, C. et al. GALNT1-mediated glycosylation and activation of Sonic hedgehog signaling maintains the self-renewal and tumor-initiating capacity of bladder cancer stem cells. Cancer Res. (2016) doi:10.1158/0008-5472.CAN-15-2309.

13. Huang, M. J. et al. Knockdown of GALNT1 suppresses malignant phenotype of hepatocellular carcinoma by suppressing EGFR signaling. Oncotarget (2015) doi:10.18632/oncotarget.3117.

14. Shan, Y. et al. LncRNA SNHG7 sponges MIR-216b to promote proliferation and liver metastasis of colorectal cancer through upregulating GALNT1. Cell Death Dis. (2018) doi:10.1038/s41419-018-0759-7.

15. Fang, R. et al. LAMTOR5 raises abnormal initiation of O-glycosylation in breast cancer metastasis via modulating GALNT1 activity. Oncogene (2020) doi:10.1038/s41388-019-1146-2.

16. Willer, C. J. et al. Newly identified loci that influence lipid concentrations and risk of coronary artery disease. Nat. Genet. (2008) doi:10.1038/ng.76.

17. Kathiresan, S. et al. Six new loci associated with blood low-density lipoprotein cholesterol, high-density lipoprotein cholesterol or triglycerides in humans. Nat. Genet. (2008) doi:10.1038/ng.75.

18. Woo, C. M., Iavarone, A. T., Spiciarich, D. R., Palaniappan, K. K. & Bertozzi, C. R. Isotope-targeted glycoproteomics (IsoTaG): A mass-independent platform for intact N- and O-glycopeptide discovery and analysis. Nat. Methods (2015) doi:10.1038/nmeth.3366.

19. Darula, Z. & Medzihradszky, K. F. Affinity enrichment and characterization of mucin core-1 type glycopeptides from bovine serum. Mol. Cell. Proteomics (2009) doi:10.1074/mcp.M900211-MCP200.

20. King, S. L. et al. Characterizing the O-glycosylation landscape of human plasma, platelets, and endothelial cells. Blood Adv. (2017) doi:10.1182/bloodadvances.2016002121.

21. Hägglund, P. et al. An enzymatic deglycosylation scheme enabling identification of core fucosylated N-glycans and O-glycosylation site mapping of human plasma proteins. J. Proteome Res. (2007) doi:10.1021/pr0700605.

22. Yang, W. et al. Comparison of Enrichment Methods for Intact N- and O-Linked Glycopeptides Using Strong Anion Exchange and Hydrophilic Interaction Liquid Chromatography. Anal. Chem. (2017) doi:10.1021/acs.analchem.7b03641.

23. Nilsson, J. et al. Enrichment of glycopeptides for glycan structure and attachment site identification. Nat. Methods (2009) doi:10.1038/nmeth.1392.

24. Halim, A., Nilsson, J., Rüetschi, U., Hesse, C. & Larson, G. Human urinary glycoproteomics; attachment site specific analysis of N- and O-linked glycosylations by CID and ECD. Mol. Cell. Proteomics (2012) doi:10.1074/mcp.M111.013649.

25. Woo, C. M. et al. Development of IsoTaG, a Chemical Glycoproteomics Technique for Profiling Intact N- and O-Glycopeptides from Whole Cell Proteomes. J. Proteome Res. (2017) doi:10.1021/acs.jproteome.6b01053.

26. Chen, M. et al. An engineered high affinity Fbs1 carbohydrate binding protein for selective capture of N-glycans and N-glycopeptides. Nat. Commun. (2017) doi:10.1038/ncomms15487.

27. Wu, A. M., Lisowska, E., Duk, M. & Yang, Z. Lectins as tools in glycoconjugate research. Glycoconjugate Journal at 10.1007/s10719-008-9119-7 (2009).

28. Yang, S. et al. Deciphering Protein O-Glycosylation: Solid-Phase Chemoenzymatic Cleavage and Enrichment. Anal. Chem. (2018) doi:10.1021/acs.analchem.8b01834.

29. Shon, D. J. et al. An enzymatic toolkit for selective proteolysis, detection, and visualization of mucin-domain glycoproteins. Proc. Natl. Acad. Sci. U. S. A. (2020) doi:10.1073/pnas.2012196117.

30. Saldova, R. & Wilkinson, H. Current methods for the characterization of o-glycans. Journal of Proteome Research at 10.1021/acs.jproteome.0c00435 (2020).

31. de las Rivas, M., Lira-Navarrete, E., Gerken, T. A. & Hurtado-Guerrero, R. Polypeptide GalNAc-Ts: from redundancy to specificity. Current Opinion in Structural Biology at 10.1016/j.sbi.2018.12.007 (2019).

32. Hintze, J. et al. Probing the contribution of individual polypeptide GalNAc-transferase isoforms to the O-glycoproteome by inducible expression in isogenic cell lines. J. Biol. Chem. (2018) doi:10.1074/jbc.RA118.004516.

33. Schjoldager, K. T. B. G. et al. Probing isoform-specific functions of polypeptide GalNAc-transferases using zinc finger nuclease glycoengineered SimpleCells. Proc. Natl. Acad. Sci. U. S. A. (2012) doi:10.1073/pnas.1203563109.

34. Vakhrushev, S. Y. et al. Enhanced mass spectrometric mapping of the human GalNAc-type O-glycoproteome with simplecells. Mol. Cell. Proteomics (2013) doi:10.1074/mcp.O112.021972.

35. Schwientek, T., Mandel, U., Roth, U., Müller, S. & Hanisch, F. G. A serial lectinapproach to the mucin-type O-glycoproteome of Drosophila melanogaster S2 cells. Proteomics (2007) doi:10.1002/pmic.200600793.

36. Gerken, T. A. et al. Emerging paradigms for the initiation of mucin-type protein O-glycosylation by the polypeptide GalNAc transferase family of glycosyltransferases. J. Biol. Chem. (2011) doi:10.1074/jbc.M111.218701.

37. Huang, J. et al. OGP: A Repository of Experimentally Characterized O-glycoproteins to Facilitate Studies on O-glycosylation. *Genomics*, Proteomics Bioinforma. (2021) doi:10.1016/j.gpb.2020.05.003.

38. Saei, A. A. et al. System-wide identification and prioritization of enzyme substrates by thermal analysis. Nat. Commun. (2021).

39. Savitski, M. M. F. et al. Tracking cancer drugs in living cells by thermal profiling of the proteome. Science (80-.). (2014) doi:10.1126/science.1255784.

40. Molina, D. M. et al. Monitoring drug target engagement in cells and tissues using the cellular thermal shift assay. Science (80-.). (2013) doi:10.1126/science.1233606.

41. Gaetani, M. et al. Proteome Integral Solubility Alteration: A High-Throughput Proteomics Assay for Target Deconvolution. J. Proteome Res. (2019) doi:10.1021/acs.jproteome.9b00500.

42. Li, J. et al. TMTpro reagents: a set of isobaric labeling mass tags enables simultaneous proteome-wide measurements across 16 samples. Nat. Methods (2020) doi:10.1038/s41592-020-0781-4.

43. Wandall, H. H. et al. Substrate specificities of three members of the human UDP-N-acetyl-α-D-galactosamine:polypeptide N-acetylgalactosaminyltransferase family, GalNAc-T1, -T2, and -T3. J. Biol. Chem. (1997) doi:10.1074/jbc.272.38.23503.

44. Uhlén, M. et al. Tissue-based map of the human proteome. Science (80-.). (2015) doi:10.1126/science.1260419.

45. Cho, N. H. et al. OpenCell: Endogenous tagging for the cartography of human cellular organization. Science (80-.). (2022) doi:10.1126/science.abi6983.

46. Tran, D. T. & Ten Hagen, K. G. Mucin-type o-glycosylation during development. Journal of Biological Chemistry at 10.1074/jbc.R112.418558 (2013).

47. Mitra, N., Sinha, S., Ramya, T. N. C. & Surolia, A. N-linked oligosaccharides as outfitters for glycoprotein folding, form and function. Trends in Biochemical Sciences at 10.1016/j.tibs.2006.01.003 (2006).

48. Lawson, E. Q. et al. Effect of carbohydrate on protein solubility. Arch. Biochem. Biophys. (1983) doi:10.1016/0003-9861(83)90449-6.

49. Huttlin, E. L. et al. Dual proteome-scale networks reveal cell-specific remodeling of the human interactome. Cell (2021) doi:10.1016/j.cell.2021.04.011.

50. Stark, C., et al. BioGRID: a general repository for interaction datasets. Nucleic Acids Res. (2006) doi:10.1093/nar/gkj109.

51. Huang, J. X. et al. High throughput discovery of functional protein modifications by Hotspot Thermal Profiling. Nat. Methods (2019) doi:10.1038/s41592-019-0499-3.

52. Orsburn, B. C. Proteome discoverer-a community enhanced data processing suite for protein informatics. Proteomes (2021) doi:10.3390/proteomes9010015.

53. Ma, B. et al. PEAKS: Powerful software for peptide de novo sequencing by tandem mass spectrometry. Rapid Commun. Mass Spectrom. (2003) doi:10.1002/rcm.1196.

54. Li, W., Hou, C., Li, Y., Wu, C. & Ma, J. HexNAcQuest: A Tool to Distinguish O-GlcNAc and O-GalNAc. J. Am. Soc. Mass Spectrom. 33, 2008–2012 (2022).

55. Sugahara, T., Pixley, M. R., Fares, F. & Boime, I. Characterization of the O-glycosylation sites in the chorionic gonadotropin β subunit in vivo using site-directed mutagenesis and gene transfer. J. Biol. Chem. (1996) doi:10.1074/jbc.271.34.20797.

56. Ou, J. et al. dagLogo: An R/Bioconductor package for identifying and visualizing differential amino acid group usage in proteomics data. PLoS One (2020) doi:10.1371/journal.pone.0242030.

57. Christlet, T. H. T. & Veluraja, K. Database analysis of O-glycosylation sites in proteins. Biophys. J. (2001) doi:10.1016/s0006-3495(01)76074-2.

58. Wilson, I. B. H., Gavel, Y. & Von Heijne, G. Amino acid distributions around O-linked glycosylation sites. Biochem. J. (1991) doi:10.1042/bj2750529.

59. O’Connell, B. C., Hagen, F. K. & Tabak, L. A. The influence of flanking sequence on the O-glycosylation of threonine in vitro. J. Biol. Chem. (1992) doi:10.1016/s0021-9258(19)73998-2.

60. Nishikawa, I. et al. Computational prediction of O-linked glycosylation sites that preferentially map on intrinsically disordered regions of extracellular proteins. Int. J. Mol. Sci. (2010) doi:10.3390/ijms11124991.

61. Trevino, S. R., Schaefer, S., Scholtz, J. M. & Pace, C. N. Increasing Protein Conformational Stability by Optimizing β-Turn Sequence. J. Mol. Biol. (2007) doi:10.1016/j.jmb.2007.07.061.

62. Potel, C. M. et al. Impact of phosphorylation on thermal stability of proteins. Nature Methods at 10.1038/s41592-021-01177-5 (2021).

63. Smith, I. R. et al. Identification of phosphosites that alter protein thermal stability. Nat. Methods (2021) doi:10.1038/s41592-021-01178-4.

64. Finn, R. D. et al. InterPro in 2017-beyond protein family and domain annotations. Nucleic Acids Res. (2017) doi:10.1093/nar/gkw1107.

65. Ward, R. A., et al. Challenges and Opportunities in Cancer Drug Resistance. Chemical Reviews at 10.1021/acs.chemrev.0c00383 (2021).

66. Medzihradszky, K. F., Kaasik, K. & Chalkley, R. J. Tissue-specific glycosylation at the glycopeptide level. Mol. Cell. Proteomics (2015) doi:10.1074/mcp.M115.050393.

67. Price, J. L. et al. Context-dependent effects of asparagine glycosylation on Pin WW folding kinetics and thermodynamics. J. Am. Chem. Soc. 132, 15359–15367 (2010).

68. Gavrilov, Y., Shental-Bechor, D., Greenblatt, H. M. & Levy, Y. Glycosylation may reduce protein thermodynamic stability by inducing a conformational distortion. J. Phys. Chem. Lett. 6, 3572–3577 (2015).

69. Hebert, D. N., Lamriben, L., Powers, E. T. & Kelly, J. W. The intrinsic and extrinsic effects of N-linked glycans on glycoproteostasis. Nat. Chem. Biol. 10, 902–910 (2014).

70. Shental-Bechor, D. & Levy, Y. Effect of glycosylation on protein folding: a close look at thermodynamic stabilization. Proc. Natl. Acad. Sci. 105, 8256–8261 (2008).

71. Sola, R. J. & Griebenow, K. Effects of glycosylate on the stability of protein pharmaceuticals. Journal of Pharmaceutical Sciences at 10.1002/jps.21504 (2009).

72. Takata, T., Oxford, J. T., Brandon, T. R. & Lampi, K. J. Deamidation Alters the Structure and Decreases the Stability of Human Lens βΑ3-Crystallin. Biochemistry 46, 8861–8871 (2007).

73. Sun, W. et al. Monitoring structural modulation of redox-sensitive proteins in cells with MS-CETSA. Redox Biol. 24, 101168 (2019).

74. Sabatier, P. et al. An integrative proteomics method identifies a regulator of translation during stem cell maintenance and differentiation. Nat. Commun. (2021) doi:10.1038/s41467-021-26879-4.

75. Saei, A. A. et al. SIESTA as a universal unbiased proteomics approach for identification and prioritization of enzyme substrates. Protoc. Exch. (2021) doi:10.21203/rs.3.pex-1327/v1.

76. Lyutvinskiy, Y., Yang, H., Rutishauser, D. & Zubarev, R. A. In silico instrumental response correction improves precision of label-free proteomics and accuracy of proteomics-based predictive models. Mol. Cell. Proteomics (2013) doi:10.1074/mcp.O112.023804.

77. Lundström, S. L. et al. Blood plasma IgG Fc glycans are significantly altered in Alzheimer’s disease and progressive mild cognitive impairment. J. Alzheimer’s Dis. (2014) doi:10.3233/JAD-131088.

78. Szklarczyk, D. et al. The STRING database in 2017: quality-controlled protein–protein association networks, made broadly accessible. Nucleic Acids Res. gkw937 (2016).

79. Vizcaíno, J. A., et al. ProteomeXchange provides globally coordinated proteomics data submission and dissemination. Nature Biotechnology at 10.1038/nbt.2839 (2014).

