## Supplementary Information for "Mapping the GALNT1 substrate landscape with versatile proteomics tools"

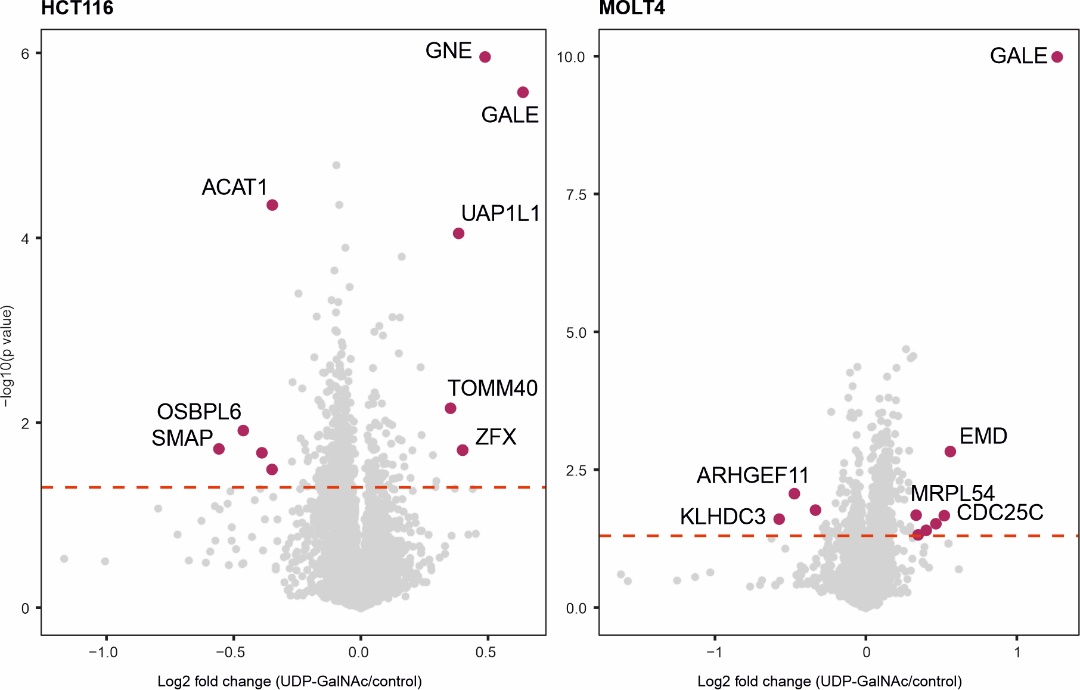

**Supplementary Fig. 1. UDP-GalNAc engages a few proteins in the cell lysate.** PISA analysis shows the engagement of few proteins including GALE and GNE in response to UDP-GalNAc at 500 μM in HCT116 and MOLT4 cell lysate.

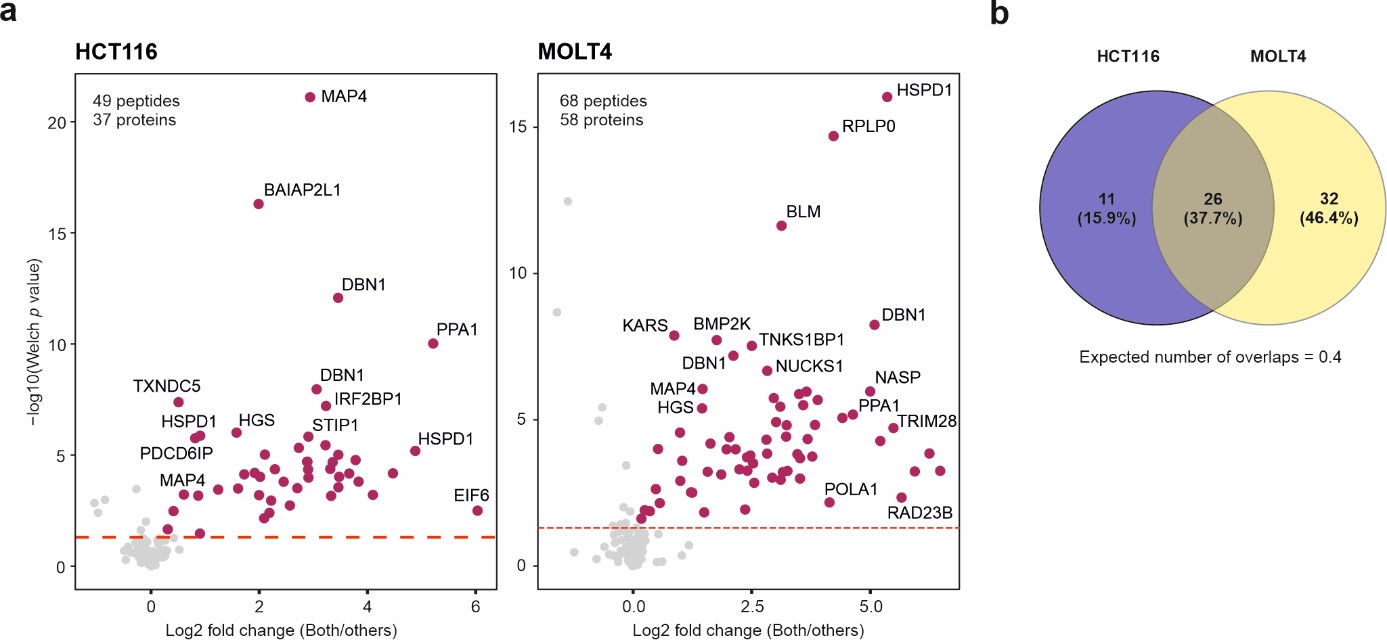

**Supplementary Fig. 2. a,** **O-HexNAc modifications identified in the SIESTA-PISA experiment.** O-glycopeptides are over-represented in the lysate treated with both GALNT1 and UDP-GalNAc compared to the other conditions, validating our experiments (two-sided Welch test; four independent biological replicates). **b,** The overlap between elevated O-glycopeptides in HCT116 vs. MOLT4 SIESTA-PISA experiments.

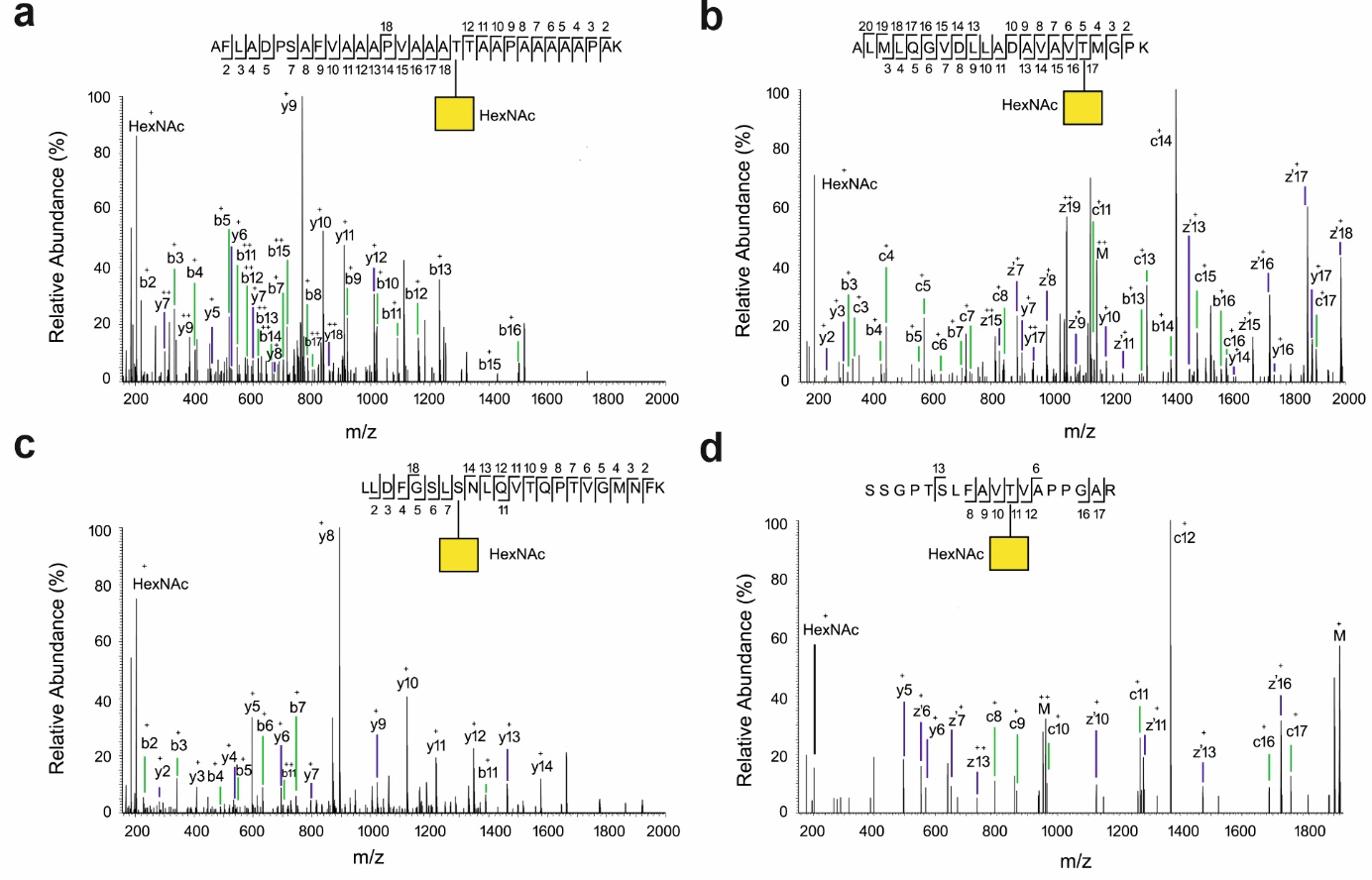
**Supplementary Fig. 3. MS/MS spectra of HexNAc modification on four peptides obtained from the glycoprotein enriched samples.** **a,** Glycopeptide from RPLP0P6 protein. **b,** Glycopeptide from CH60 protein. **c**, Glycopeptide from RPS17 protein. **d**, Glycopeptide from HNRPU protein. All four glycopeptides are shared between the two cell lines and significant between the lysate treated with both GALNT1 and UDP-GalNAc compared to the control (**Supplementary data 6**). Note that the suggested position assignment of the glycan, particularly for a and c (obtained from HCD- spectra), is not ambiguous.

**Supplementary Table 1.** The number of assigned GlcNAc and GalNAc isomers in the glycoprotein enriched samples using HexNAcQuest search tool. The search results with the diagnostic oxonium ions are given in **Supplementary Data 7**.

|  |  | GlcNAc | GalNAc | GalNAc/GlcNAc | log2(GalNAc/GlcNAc) |
| --- | --- | --- | --- | --- | --- |
| MOLT4 | Both1 | 91 | 125 | 1.374 | 0.46 |
|  | Both2 | 28 | 61 | 2.179 | 1.12 |
|  | Both3 | 61 | 133 | 2.180 | 1.12 |
|  | C1 | 59 | 9 | 0.153 | -2.71 |
|  | C2 | 60 | 8 | 0.133 | -2.91 |
|  | C3 | 58 | 7 | 0.121 | -3.05 |
| HCT116 | Both1 | 53 | 82 | 1.547 | 0.63 |
|  | Both2 | 55 | 100 | 1.818 | 0.86 |
|  | Both3 | 53 | 76 | 1.434 | 0.52 |
|  | C1 | 32 | 7 | 0.219 | -2.19 |
|  | C2 | 31 | 5 | 0.161 | -2.63 |
|  | C3 | 30 | 5 | 0.167 | -2.58 |

**Supplementary Table 2.** The enriched domains in the substrate proteins

| **Domain** | **Description** | **Count** | **Strength** | **FDR** |
| --- | --- | --- | --- | --- |
| IPR024084 | Isopropylmalate dehydrogenase-like domain | 4 of 5 | 1.37 | 0.043 |
| IPR019818 | Isocitrate/isopropylmalate dehydrogenase, conserved site | 4 of 5 | 1.37 | 0.043 |
| IPR000558 | Histone H2B | 12 of 17 | 1.32 | 6e-08 |
| IPR007125 | Histone H2A/H2B/H3 | 17 of 43 | 1.07 | 9e-09 |
| IPR004001 | Actin, conserved site | 6 of 16 | 1.04 | 0.024 |
| IPR037103 | Tubulin/FtsZ, C-terminal domain superfamily | 7 of 21 | 0.99 | 0.018 |
| IPR020902 | Actin/actin-like conserved site | 6 of 18 | 0.99 | 0.036 |
| IPR023313 | Ubiquitin-conjugating enzyme, active site | 8 of 25 | 0.97 | 0.007 |
| IPR018316 | Tubulin/FtsZ, 2-layer sandwich domain | 7 of 22 | 0.97 | 0.020 |
| IPR023123 | Tubulin, C-terminal | 7 of 23 | 0.95 | 0.024 |
| IPR017975 | Tubulin, conserved site | 7 of 23 | 0.95 | 0.024 |
| IPR008280 | Tubulin/FtsZ, C-terminal | 7 of 23 | 0.95 | 0.024 |
| IPR003008 | Tubulin/FtsZ, GTPase domain | 7 of 23 | 0.95 | 0.024 |
| IPR000217 | Tubulin | 7 of 23 | 0.95 | 0.024 |
| IPR036525 | Tubulin/FtsZ, GTPase domain superfamily | 7 of 24 | 0.93 | 0.024 |
| IPR009072 | Histone-fold | 17 of 72 | 0.84 | 5e-06 |
| IPR000608 | Ubiquitin-conjugating enzyme E2 | 9 of 38 | 0.84 | 0.014 |
| IPR016135 | Ubiquitin-conjugating enzyme/RWD-like | 11 of 53 | 0.79 | 0.007 |
| IPR035979 | RNA-binding domain superfamily | 45 of 241 | 0.74 | 1e-14 |
| IPR000504 | RNA recognition motif domain | 41 of 227 | 0.73 | 6e-13 |
| IPR012677 | Nucleotide-binding alpha-beta plait domain superfamily | 42 of 242 | 0.71 | 6e-13 |

Supplementary Data Files:

**Supplementary Data 1.** SIESTA-PISA analysis in HCT116 and MOLT4 cell extracts

**Supplementary Data 2.** The compiled list of substrates with further annotations

**Supplementary Data 3.** Proteins interacting with GALNT1 in HCT116 and MOLT4 cell extracts (in PISA analysis)

**Supplementary Data 4.** Peptides carrying HexNAc modification in SIESTA-PISA data, and their occupancies

**Supplementary Data 5.** Glycoprotein enrichment data for HCT116 and MOLT4

**Supplementary Data 6.** Peptides carrying HexNAc modification in glycoprotein enrichment data

**Supplementary Data 7**. O-GlcNAc vs O-GalNAc search analysis using HexNAcQuest
